# Case report of unusual mortality of Egyptian fruit bat (*Rousettus aegyptiacus*) in northern Cyprus

**DOI:** 10.64898/2026.06.08.730983

**Authors:** Maya Weinberg, Dominika Knazovicka, Rianne Pelgrims, Ilayda Taskaya, Pera Sinkovec, Luis Víquez-R, Kendra Phelps, Allyson Walsh, Paul A. Racey, Tigga Kingston, Julie Teresa Shapiro

## Abstract

We report an unusual mortality event affecting the isolated Egyptian fruit bat (*Rousettus aegyptiacus*) population in northern Cyprus, combining clinical admissions, microbiological findings, and roost surveys to assess magnitude and potential drivers. From January to June 2025, the Cyprus Wildlife Research Institute (CWRI) Wildlife Hospital received an unprecedented surge in admissions with 12 individuals (versus sporadic admissions in prior years), with high acute mortality: most bats arrived moribund and died within 12–24 hours.

Clinical records noted localized purulent lesions and abscesses in a substantial fraction of cases. Bacteriological culture and PCR assays recovered *Staphylococcus aureus* from abscesses and skin swab samples in multiple individuals; isolates exhibited susceptibility to tested antibiotics. Necropsy sampling confirmed viable *S. aureus* in several specimens despite prolonged storage. Concurrent targeted surveys of ten known roosts in Feb–Mar 2026 documented marked declines at multiple historical sites, including reductions from hundreds to single-digit counts at formerly large colonies and the complete absence of bats at multiple roosts. No clear evidence of recent human disturbance, extreme weather anomalies, or reduced food availability was found; shotgun cartridges from historical hunting were present, but no direct anthropogenic cause was apparent. While *S. aureus* infections—seasonally concentrated in winter—are consistent with observed lesions and may have contributed to morbidity and mortality, causality for the population-level declines remains unknown, and other factors (toxins, unassessed pathogens, multi-factor stressors) cannot be excluded. Given the genetic isolation and conservation significance of the Cyprus Egyptian fruit bat population, these findings are concerning. We recommend urgent, coordinated longitudinal population monitoring, expanded pathogen surveillance including whole-genome sequencing of *S. aureus* isolates, toxicological screening, and development of a species recovery plan incorporating emergency response, habitat protection, and public outreach.

## Introduction

The Egyptian fruit bat (*Rousettus aegyptiacus*, Geoffroy, 1810) is a species belonging to the family Pteropodidae. It has a widespread distribution from southern Africa, from Senegal in the west up to Egypt, the Arabian Peninsula, Levant, Türkiye, and Cyprus, and west to Pakistan (Bergmans 1994; Kwiecinski and Griffiths 1999; Korine 2016; Strachinis et al. 2018; Monadjem et al. 2024). The island of Cyprus is divided into the Republic of Cyprus in the south and the self-declared Turkish Republic of Northern Cyprus in the north—recognized only by Türkiye.

The island is home to the European Union’s only population of the Egyptian fruit bat, or any pteropodid species. This island population appears isolated from the closest mainland populations in Türkiye and is thus genetically distinct (Hulva et al. 2012).

Egyptian fruit bats face multiple anthropogenic threats in northern Cyprus. Starting in the 1920s, the British colonial government on the whole island carried out extermination campaigns, which were halted in northern Cyprus when it came under Turkish military control in 1974 (Hadjisterkotis 2006). In the Republic of Cyprus, sudden, precipitous declines in the population numbers have been observed, although the cause remains unknown (Nicolau 2009; del Vaglio et al. 2011). While population data from northern Cyprus is more sparse than from the Republic of Cyprus (Baydemir 2014), a major decline was observed from 2010 to 2018, with multiple roosts reduced from hundreds of individuals to mere tens and the complete abandonment of several previously occupied sites (Benda et al. 2018).

Bacterial disease appears to be another overlooked but potentially growing threat to bats (Evans et al. 2020; Imnadze et al. 2020), including the Egyptian fruit bat (Weinberg et al. 2022). *Staphylococcus aureus* is a common opportunistic pathogenic bacterium, although individuals may also be carriers with no signs of disease (Fountain et al. 2022). It typically acts as a secondary opportunistic pathogen, with clinical disease often developing in hosts already compromised by other primary infections or by environmental or physiological conditions that favor bacterial proliferation (Haag et al. 2019). Nevertheless, the social structure and frequent interactions within bat colonies can facilitate rapid spread between individuals (Razik et al. 2026). Previous studies have detected *S. aureus* in several bat species worldwide (Akobi et al. 2012; Olatimehin et al. 2018; Attaullah et al. 2026), including in Egyptian fruit bats (Held et al. 2017; Ngoubangoye et al. 2026). In Israel, bacterial disease in free-ranging Egyptian fruit bats presenting severe skin lesions from which *S. aureus* was consistently isolated has been documented as a major source of morbidity (Weinberg et al. 2022).

Here, we describe an apparent mortality event among Egyptian fruit bats in northern Cyprus in 2025, evidenced by a marked increase in admissions to a local wildlife hospital, including individuals presenting purulent lesions for which we present bacterial findings. Simultaneously, we observed a generalized population decline in 2025, in some cases quite steep, at known roosts in the region, including caves, mines, quarries, and man-made structures. We present current counts at these roosts and compare them with estimates from previous observations.

### Case Description

#### Admissions and case presentations

Medical admissions of Egyptian fruit bats to the Wildlife Hospital of the Cyprus Wildlife Research Institute, a local non-governmental, non-profit nature conservation and research organization based in Taşkent, Kyrenia (northern Cyprus), were rare before the study period (data collected since 2016). Records indicate that one individual was admitted to the hospital in each of the years 2017, 2018, three in 2019, and two in 2020, while no bats were recorded in 2021 or 2022, followed by five individuals in 2023 and four in 2024. None of the cases admitted to the hospital before 2025 exhibited any observable abscesses or symptoms of a clear bacterial disease.

In 2025, 12 Egyptian fruit bats were submitted to the Wildlife Hospital (Table 1). Admissions occurred between January and June 2025, with a clear peak in late winter, when four bats were admitted in February and six in March, representing the highest concentration of cases over the year. The bats originated from several regions across northern Cyprus, including Lefkoşa (Nicosia; n = 5), Girne (Kyrenia; n = 6), and Gazimağusa (Famagusta; n = 1) (Table 1; Figure 1). As of this writing (May 2026), no Egyptian fruit bats have been admitted in 2026.

**Table 1.** Characteristics of the 12 Egyptian fruit bats (*Rousettus aegyptiacus*) brought to the Wildlife Hospital of the Cyprus Wildlife Research Institute in 2025 that were screened for bacterial infections. Two additional Egyptian fruit bats admitted in 2020 and 2024 and two Kuhl’s pipistrelles (*Pipistrellus kuhlii*) admitted in 2025 that were sampled for bacterial analysis in February 2026 are also included.

| Species | Bat ID | Admission date<br>(mm/dd/yyyy) | Region | Sex* | Age** | Body mass<br>(g) | Outcome*** | Abscess /<br>lesion | Bite wounds | Bacterial screening<br>samples<br>(year) | *Staphylococcus*<br>*aureus* isolate<sup>+</sup><br>(year) |
| --- | --- | --- | --- | --- | --- | --- | --- | --- | --- | --- | --- |
| *Rousettus*<br>*aegyptiacus* | RC1467 | Jul. 11 2020 | Kyrenia | F | A | 79 | DOA | No | No | Yes (2026) | No |
|  | RC4617 | Nov. 25 2024 | Nicosia | M | A | 108 | DOA | No | No | Yes (2026) | Yes<sup>+</sup> (2026) |
|  | RC4642 | Jan. 1 2025 | Kyrenia | F | A | 129 | DOA | No | Yes | No | No |
|  | RC4660 | Feb. 4 2025 | Nicosia | M | A | 149 | DOA | No | Yes | No | No |
|  | RC4665 | Feb. 13 2025 | Nicosia | F | A | 132 | DOA | Yes | No | Yes (2026) | Yes |
|  | EXTD0520 | Feb. 14 2025 | Nicosia | M | A | 141 | Carcass | No | No | No | No |
|  | RC4667 | Feb. 16 2025 | Nicosia | M | A | 142 | D24H | Yes | Yes | Yes (2025, 2026) | Yes<sup>+</sup> (2025, 2026) |
|  | RC4681 | Mar. 8 2025 | Kyrenia | F | A | 118 | DBE | Yes | Yes | Yes (2025, 2026) | Yes (2025) |
|  | EXTD0521 | Mar. 15 2025 | Nicosia | M | A | 136 | Carcass | Yes | Yes | Yes (2025, 2026) | Yes (2025) |
|  | EXTD0524 | Mar. 17 2025 | Kyrenia | N/A | A | N/A | Carcass | N/A | No | Yes (2026) | No |
|  | RC4690 | Mar. 17 2025 | Kyrenia | F | A | 108 | DOA | Yes | Yes | Yes (2025, 2026) | Yes (2025, 2026) |
|  | RC4691 | Mar. 20 2025 | Famagusta | F | A | 117 | D24H | Yes | Yes | Yes (2026) | No |
|  | EXTD0526 | Mar. 26 2025 | Kyrenia | F | A | 112 | Carcass | No | Yes | Yes (2026) | No |
|  | RC4930 | Jun. 25 2025 | Kyrenia | F | A | 110 | DOA | Yes | No | Yes (2026) | Yes<sup>+</sup> (2026) |
| *Pipistrellus*<br>*kuhlii* | RC4871 | May 29 2025 | NA | M | J | 1.1 | EUE | No | No | Yes (2026) | No |
|  | RC5021 | Sep. 16 2025 | NA | N/A | Adult | 3.0 | DOA* | No | No | Yes (2026) | No |

\*Sex: F - Female; M - Male
\*\*Age: A - Adult; J - Juvenile
\*\*\*Outcome categories: DOA - dead on arrival; DBE - died before examination; D24H - dead within 24 hours after admission; EUE - euthanized upon examination; Carcass - received as decomposing carcass
<sup>+</sup>Indicates *Staphylococcus aureus* identification of isolate confirmed by whole genome sequencing (2026 isolates only)

**Figure 1.**
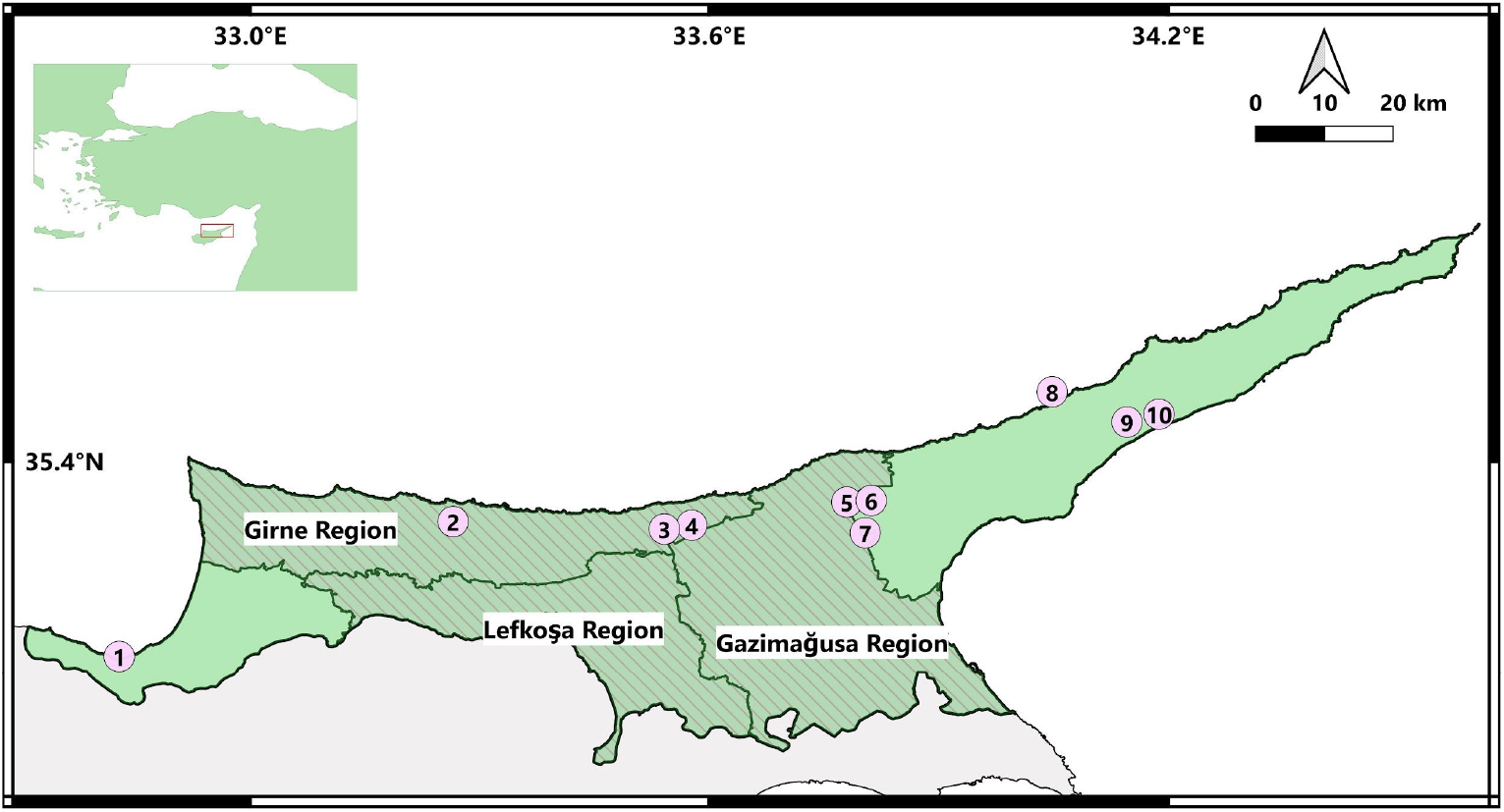
Map of northern Cyprus (light green) showing the regions of origin of the Egyptian fruit bats submitted to the Wildlife Hospital in 2025 (shaded) and locations of roosts visited during field surveys in February to March 2026 (Table 2). 1. Bağlıköy, 2. Ağırdağ, 3. Alevkaya Canyon, 4. Görneç, 5. İncirli, 6. Altınova site 2, 7. Altınova site 1, 8. Yedikonuk, 9. Kumyalı site 1, and 10. Kumyalı site 2.

Of the 12 admitted Egyptian fruit bats, four were deceased at the time of initial reporting via a call to the rescue hotline, five died before arrival at the Wildlife Hospital, and the remaining three died within 24 hours of admission. Seven were females and four were males, while the sex of one individual could not be determined. The average body mass of males was 142 g, whereas females averaged 118 g, which aligns with the normal body weight range and sexual dimorphism between males and females of the species in this region (Albayrak et al. 2008). All 12 individuals were classified as adults in the clinical records based on their size and by examining ossification of the epiphyseal joints by shining a light behind a finger joint in the wing (Racey 1974). All bats had empty stomachs on arrival, which is typical for this species given their rapid digestion (Docters van Leeuwen 1935; Tedman and Hall 1985; Shilton et al. 1999) and not indicative of pathology. Holes in the patagium were observed in three individuals, and visible wounds (resembling bite wounds) were observed on the torso and wings of eight individuals. Bite wounds were assessed as peri-mortem in five cases and possibly post-mortem in three cases, likely inflicted by free-ranging or feral cats (Oedin et al. 2021), which are widespread in the area. In all cases, the wounds were minor and were not considered a likely or significant contributor to death in any of the bats, although they could indicate a weakened state, allowing the bats to be predated by cats. Abscesses were recorded in seven bats, indicating the presence of localized purulent lesions (Table 1; Figure 2).

**Table 2.** Overview of historical and recent visits to Egyptian fruit bat roost sites in northern Cyprus.

| Site name | Date of visit | Count | Source |
| --- | --- | --- | --- |
| Bağlıköy | November 2022 | 500 | CWRI (unpublished data) |
|  | March 2023 | 500 | CWRI (unpublished data) |
|  | August 2024 | 500 | CWRI (unpublished data) |
|  | March 2025 | 4 | CWRI (unpublished data) |
|  | February 2026 | 5 | (this study) |
| Görneç | January 2018 | 30 | Benda et al. 2018 |
|  | May 2018 | 20 | Benda et al. 2018 |
|  | October 2018 | 40 | Benda et al. 2018 |
|  | February 2026 | 40 | (this study) |
| Kumyalı site 1 | May 2018 | feeding roost | Benda et al. 2018 |
|  | November 2022 | 0 | CWRI (unpublished data) |
|  | February 2026 | 0 | (this study) |
| Kumyalı site 2 | May 2018 | 20 | Benda et al. 2018 |
|  | October 2018 | 6 | Benda et al. 2018 |
|  | February 2026 | 0 | (this study) |
| Yedikonuk | October 2005 | 800 | Benda et al. 2007, Hulva et al. 2012 |
|  | May 2009 | 2 | Benda et al. 2011 |
|  | February 2010 | 20 | Benda et al. 2011 |
|  | November 2015 | 40 | Benda et al. 2018 |
|  | October 2018 | 20 | Benda et al. 2018 |
|  | March 2025 | 1 | CWRI (unpublished data) |
|  | February 2026 | 15 | (this study) |
| Alevkaya Canyon | July 2015 | Detected calls | Benda et al. 2018 |
|  | August 2016 | Detected calls | Benda et al. 2018 |
|  | May 2018 | Detected calls | Benda et al. 2018 |
|  | February 2026 | No calls detected | (this study) |
| Ağırdağ | April 2005 | 12 | Benda et al. 2007 |
|  | May 2009 | 2 | Benda et al. 2018 |
|  | August 2009 | 5 | Benda et al. 2011 |
|  | March 2026 | 0 | (this study) |
| Altınova site 1 | January 2018 | 1 | Benda et al. 2018 |
|  | March 2026 | 0 | (this study) |
| Altınova site 2 | January 2018 | 20 | Benda et al. 2018 |
|  | March 2026 | 20 | (this study) |
| İncirli | June 2002 | 5 | Anecdotal reports |
|  | November 2022 | 0 | CWRI, unpublished data |
|  | February 2026 | 0 | (this study) |

**Figure 2.**
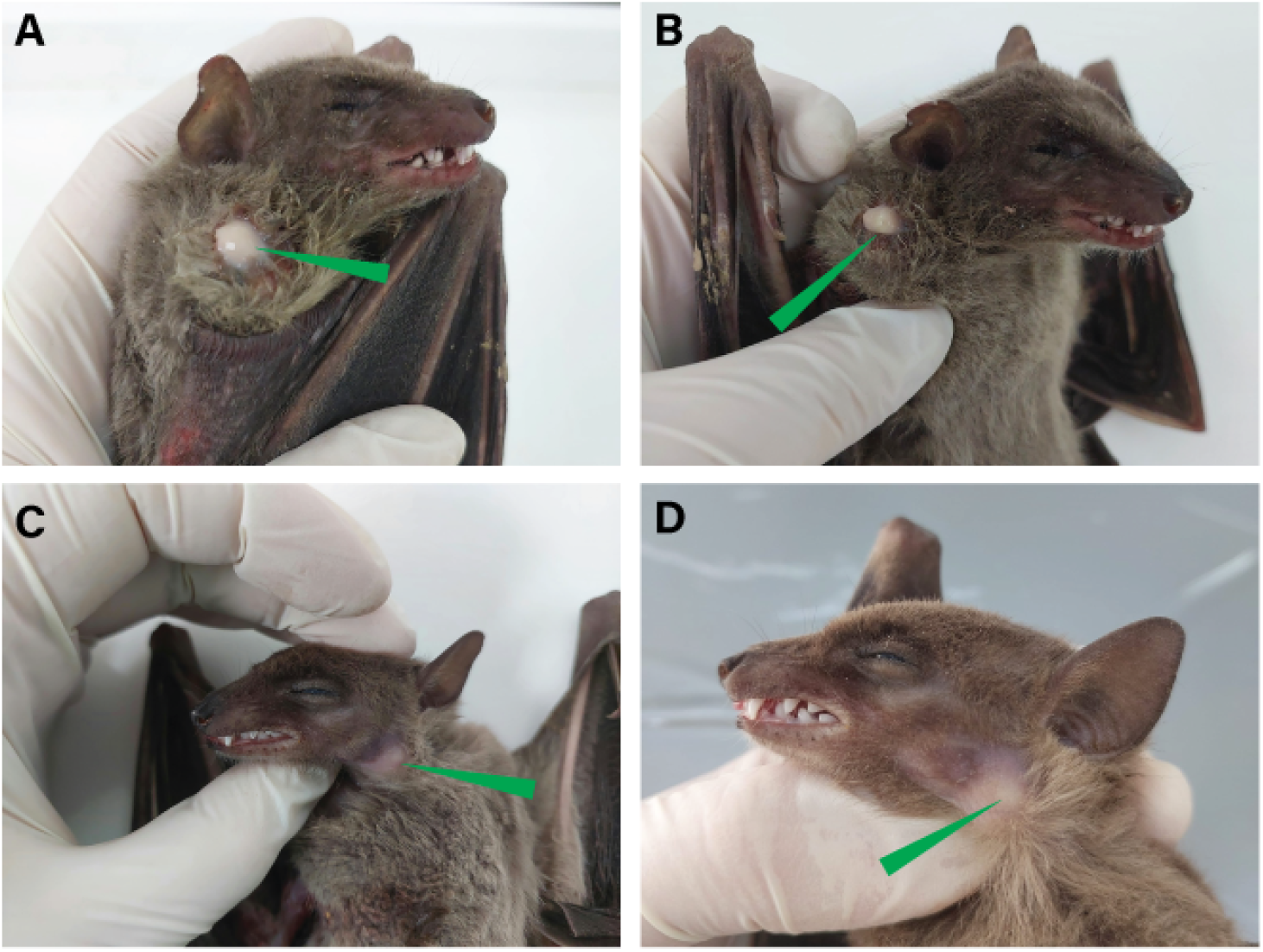
(A–B) Individual (Bat ID RC4667) presented with an abscess on the right shoulder; (C–D) Individual (Bat ID RC4691) presented with an abscess on the left side of the upper neck. These abscesses are the typical clinical appearance of bacterial infection with purulent (pus-filled) content. Photographs provided by Rianne Pelgrims, March 2025.

During the peak admission period (January - March 2025), purulent discharge was drawn from cervical abscesses of four bats for bacterial culture on discriminant plates and antimicrobial susceptibility testing via minimum inhibitory concentration (MIC) using the VITEK 2 system (bioMérieux) (Ligozzi et al. 2002) (Table 1). The cultured bacteria were identified as *S. aureus* via visual observation of cultures, the catalase test (Koneman et al. 1992), and identification with the VITEK 2 system (Ligozzi et al. 2002). No detectable antimicrobial resistance was found. The isolates were not retained and could therefore not be further characterized. All other bacteriological analyses were carried out in 2026 on frozen bat carcasses.

#### Bacteriological analyses

The Wildlife Hospital maintains bat carcasses frozen at −20°C at their facilities. On February 18, 2026, a total of 11 Egyptian fruit bats were thawed for examination and sampling for bacteriological analyses. Nine of these bats were admitted to the hospital in 2025, one in 2020, and one in 2024 (Table 1). For the bats admitted in 2025 (n=9), four of them had already been sampled for bacterial culture and antimicrobial susceptibility testing in 2025 and yielded isolates as described above. However, because these isolates had not been preserved, these individuals were sampled again in 2026.

The sampling in February 2026 involved taking fur and skin swabs and swabs from abscesses or lesions. Tissue or cavity samples from internal organs such as the lung, heart, liver, kidney, spleen, and intestinal tract were also taken for future virological and fungal analyses. In addition, the carcasses of two Kuhl’s pipistrelles (*Pipistrellus kuhlii*), an insectivorous verspertilionid species, admitted in 2025, were sampled for both bacteriological analyses and future virological and fungal analyses as an interspecies comparison (Table 1).

To culture and isolate any bacteria in the lesions, all swabs were inoculated into 10 mL of 6% NaCl enrichment broth and incubated statically at 37°C for 24 hours. Our protocol targeted *S. aureus* because the appearance and location of lesions and abscesses were similar to those previously observed in infections caused by this bacterial species in Egyptian fruit bats in Israel (Weinberg et al. 2022). Cultures were then mixed to homogenize them in the liquid, and aliquots (ranging from 5-10 µL, depending on optical density) were plated onto Brilliance™ Staph 24 agar discriminative plates and incubated at 37°C for 24 hours. Single colonies from positive plates were subcultured onto blood agar and incubated under the same conditions to ensure purity. From the blood agar plates, pure isolates were taken for molecular analysis.

In total, eight pure isolates originating from five individuals were recovered from the swabs and identified as *S. aureus*. Of these five individual bats, one was admitted in 2024 without any abscesses or lesions; the other four were admitted in 2025 and all presented with abscesses or lesions (Table 1). To confirm *S. aureus* identity, a PCR assay targeting the femB gene was performed on pure isolates. Colonies confirmed by PCR were then inoculated into 5 mL tryptic soy broth (TSB) and incubated overnight at 37°C. From the overnight incubation of pure isolates, 1 mL of culture was centrifuged at maximum speed for 3 minutes to obtain a bacterial pellet. This pellet was either used for DNA extraction using the MasterPure™ Gram Positive DNA Purification Kit (Lucigen) or shipped as-is to MicrobesNG (Birmingham, UK) for DNA extraction and subsequent genomic analysis.

Three pure isolates, each from a different individual bat, were sent for whole-genome sequencing (WGS) (Table 1) with results pending.

#### Population sampling

To evaluate potential population declines concurrent with the reported increase in mortalities, we surveyed 10 known roosting sites of Egyptian fruit bats in February and March 2026. At each roost, we counted the bats present and compared their abundance with historical observations (Table 2). We noted a decline in the number of bats at all but one of these sites. Declines were particularly dramatic at Bağlıköy dropping from an estimated 500 individuals to five between 2024 and 2026. Similarly, at Yedikonuk, the number of roosting bats has declined from 800 in 2005 to 15 in 2026. At five sites where Egyptian fruit bats or signs of their presence (e.g., feces and uneaten fruit) had been previously observed, no individuals or other signs of roost occupation could be found.

## Discussion

We report a marked, recent decline of Egyptian fruit bats across northern Cyprus, documented by dramatic reductions at multiple historical roosts and an unusual surge in clinical admissions to the Cyprus Wildlife Research Institute in 2025. Although the IUCN Red List classifies the Egyptian fruit bat as a species of “Least Concern” with a globally stable population (Korine 2016), its population on Cyprus is small, vulnerable, and genetically isolated (Hulva et al. 2012; Benda et al. 2012). The mortality event and the significant population decline across known roosts that we observed in northern Cyprus are cause for conservation concern.

While the 12 admissions to the CWRI Wildlife Hospital may not appear extreme, it is, in fact, highly concerning and unusual given how generally rare it is for members of the public to find bats or their carcasses. In fact, none of the bats were actively collected from natural roosts or through targeted sampling; all of these bats were found by members of the public and either brought directly to the CWRI Wildlife Hospital or transported there by the rescue team. In most years, no Egyptian fruit bats are brought in, and the highest previous count was five individuals. Generally, bat cadavers are rarely found by the public since most die in inaccessible locations or are either quickly scavenged or decompose (Arnett et al. 2008; Bastos et al. 2013). In this context, and combined with the steep declines at known roost sites, the admissions likely represent only a very small fraction of the total mortalities in this Egyptian fruit bat population.

The precise cause – or causes – of the observed mortality and population declines remain unknown. Clinical records and necropsy sampling from the Egyptian fruit bats brought to the CWRI Wildlife Hospital revealed frequent localized purulent skin lesions and abscesses located primarily in the neck and upper back. This is consistent with previously documented *S. aureus* infections in Egyptian fruit bats in Israel during the winter months, coinciding with the period of highest physiological stress (Weinberg et al. 2022). Although Weinberg et al. (2022) did not report mortality associated with the infections, it is plausible that severe cases could progress to death if followed longitudinally. Further, the isolation of *S. aureus* from abscesses, which are not optimal for long-term bacterial preservation, suggests a substantial bacterial burden among the affected individuals. While it is generally thought that *S. aureus* infections contribute to morbidity as an opportunistic pathogen but are unlikely to be the root cause of mortality (instead indicating a generally weakened state due to other factors) (Weinberg et al. 2022), severe *S. aureus* infections have been observed to cause death or debilitating injuries that impede flight in vampire bats in captivity (Razik et al. 2026). At this time, we are unable to rule out these infections as a possible cause of death.

More generally, the winter peak in admissions, adult-biased cases, lack of clear human disturbance at roosts, and typical environmental conditions, food availability, and body mass of admitted bats argue for a biologically driven event rather than an acute, localized human action. However, other factors (e.g., toxins, unmeasured pathogens, and nutritional stress) cannot be excluded. Reports from roosts in both northern and southern Cyprus (Dechmann et al. 2026), including observations shared on social media, suggest a similar peak in mortality during the same winter period across the island, indicating a potentially comparable pattern across the island.

Given the uncertainties regarding the mortality event and population decline of Egyptian fruit bats in northern Cyprus, urgent, coordinated investigations and monitoring are needed to elucidate the cause of this mortality event and inform effective countermeasures (Figure 3). Future work in the region should adopt an integrated, longitudinal framework that combines disease surveillance, ecological monitoring, and applied conservation. Pathogen surveillance should be expanded through systematic bacteriological, viral, and fungal screening of live, moribund, and freshly dead bats, with particular emphasis on whole-genome sequencing of *S. aureus* isolates to compare strains from healthy and symptomatic individuals and across neighboring regions. In parallel, coordinated follow-up of population dynamics using standardized, repeated roost counts, alongside mark–recapture studies, would enable robust estimates of population size, demographic structure, reproductive success, and seasonal trends. These datasets should be leveraged to develop risk-assessment models that project disease spread and population trajectories under alternative scenarios, including pathogen-driven mortality and multi-factor stressors. Such analyses would help identify high-risk roosts and critical periods, estimate local extinction risk, and evaluate the potential for population rescue from adjacent areas. Ultimately, this knowledge should inform the development of a species recovery plan incorporating emergency response protocols (e.g., rescue, treatment, and carcass handling), targeted habitat protection and disturbance minimization at key roosts, and public outreach initiatives aimed at reducing anthropogenic pressures and supporting long-term population recovery.

**Figure 3.**
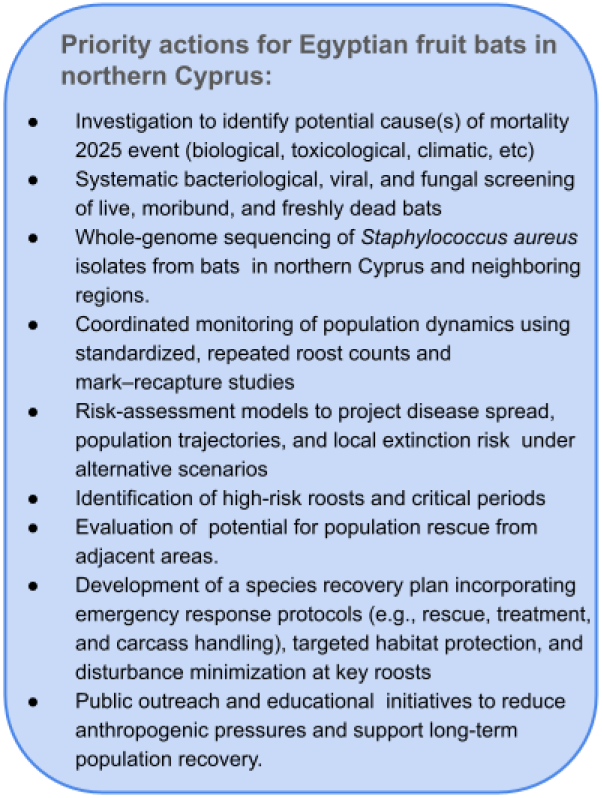
Priority action items for conservation of the Egyptian fruit bat (*Rousettus aegyptiacus*) in northern Cyprus.

## Data availability

All data associated with this study are presented in the main text of this case report.

## Conflict of interest

The authors declare that there is no conflict of interest regarding the publication of this article

## Funding statement

This work was not funded by any specific grants or funding.

## Acknowledgements

We gratefully acknowledge Enzo Moraes and Iona Shearer (CWRI) for their valuable logistical support during the fieldwork. We further thank Kemal Basat for his financial support of CWRI’s work.

## Notes

### Competing Interest Statement

The authors have declared no competing interest.

